# A duration-dependent interaction between high-intensity light and optical refocus in the drive for myopia control

**DOI:** 10.1101/2022.12.19.520946

**Authors:** Sayantan Biswas, Arumugam R. Muralidharan, Bjorn Kaijun Betzler, Joanna Marie Fianza Busoy, Veluchamy A. Barathi, Royston K. Y. Tan, Wan Yu Shermaine Low, Dan Milea, Biten K. Kathrani, Noel A. Brennan, Raymond P. Najjar

## Abstract

**PURPOSE:** To evaluate the duration-dependent and synergetic impact of high-intensity light (HL) and optical refocus (RF) on lens-induced myopia (LIM) development in chickens.

**METHODS:** Myopia was induced in one eye in chicks (10 groups, n=126) from day 1 post-hatching (D1) until D8 using -10D lenses. Fellow eyes remained uncovered as controls. Nine groups were exposed daily to continuous 2 hours (h), 4h, or 6h of either HL (15,000 lux); RF (removal of -10D lens); or both (HL+RF). One group served as the LIM group without any interventions. Ocular axial length (AL), refractive error, and choroidal thickness were measured on D1, D4, and D8. Outcome measures are expressed as inter-ocular difference (IOD= experimental - control eye) ±SEM.

**RESULTS:** By D8, LIM increased AL (0.36±0.04 mm), myopic refraction (-9.02±0.37D), and choroidal thinning (-90.27±16.44 µm) in the LIM group (all, P<0.001). Compared to the LIM group, exposure to 2h, 4h, or 6h of HL, RF, or HL+RF reduced myopic refraction in a duration-dependent manner, with RF being more effective than HL (P<0.05). Only 6h of HL+RF (not 2h or 4h) prevented LIM and was more effective than RF (P=0.004) or HL (P<0.001) in reducing myopic refraction, and more effective than HL (P<0.001) in reducing axial elongation.

**CONCLUSION:** Daily exposure to 2h, 4h, or 6h of HL, RF, or HL+RF reduced lens-induced myopic refraction in a duration-dependent manner in chickens. Only 6h of HL+RF completely stopped LIM development. The synergetic effect of HL and RF is dependent on the duration of the interventions.

## Introduction

Myopia, also known as near-sightedness, is one of the most prevalent eye conditions worldwide and has become a public health concern, particularly in East and Southeast Asia.^1^ Axial myopia, the most common form of myopia, involves excessive globe elongation during ocular development, resulting in a mismatch between the axial length (AL) and optical power of the eye.^2^ While the refractive error associated with myopia can be corrected, high myopia is associated with sight-threatening pathologies, including retinal detachment, glaucoma and myopic maculopathy.^3^

Environmental factors alongside genetics are known to play a significant role in the development of myopia.^4^ Increased outdoor activity in childhood reduces the risk of developing myopia,^5–7^ independent of physical activity.^8^ This protective impact of outdoor time may be due to a combination of factors, including the intensity and spectral composition of sunlight,^9^ the increased spatial frequency and higher image contrast encountered outdoors,^10,11^ and the significant differences in the pattern of retinal defocus generated outdoors compared with indoors,^10,12^ associations that are still under investigation in animal models of myopia.^13^ Because time spent outdoors time involves all of these factors simultaneously, the relative protective contribution of each and interaction between them is unknown. Animal models offer an opportunity to explore such associations.

The most commonly used myopia induction methods in animal models are form-deprivation myopia (FDM), in which translucent diffusers are fitted to obscure the animal’s vision, and lens-induced myopia (LIM), in which hyperopic defocusing lenses (i.e., negative powered lenses) are fitted to place the focal plane behind the retina and induce axial ocular growth.^14^ FDM and LIM have overlapping, yet different underlying mechanisms of action^15,16^ – caution is required when extrapolating the results of each experimental model to the other.^15^ For instance, FDM and LIM both involve lowered retinal dopamine levels, but growth control mechanisms and response to atropine differ between the two.^16,17^

Exposure to high-intensity light (HL, 10,000-25,000 lux/day) can be effective in alleviating FDM in various animal models.^18–21^ In addition, Ashby et al.^22^ reported that in chicks, the suppression of FDM is even more pronounced when HL was combined with normal vision (diffuser removal) for 15 minutes/day, compared to experimental groups exposed to either HL (15,000 lux) or diffuser removal alone. The authors postulated that dopamine release in the chick retina had a graded response to increasing illumination due to diffuser removal, resulting in greater inhibition of axial ocular growth at higher illumination levels.^22^ Another hypothesis was that higher illuminations led to greater pupil constriction, greater depth of focus, and reduced image blur, promoting the developing eye’s attempts at maintaining emmetropia. In studies utilizing LIM, exposure to HL without the defocusing lens removal or optical refocus (RF) significantly slowed LIM development in chicks (15,000 lux for 5 hours (h)/day),^23^ but not in monkeys (25,000 lux for 6h/day),^24^ while exposure to brief periods (1-3h) of RF/day reduced the extent of LIM development in tree shrews,^25,26^ marmosets,^27^ and chicks.^28^ While LIM and FDM may yield different ocular mechanisms,^15,16^ lens removal for LIM and normal vision in case of FDM may operate through different pathways. In LIM, RF may act through a change in optical and visual spatial information, whereas normal vision in FDM additionally involve neuromodulations associated with increased retinal illumination.^29–31^

To our knowledge, the dose-dependent and synergetic effects of HL, RF and HL+RF are yet to be investigated in LIM models. This study investigates the interactive impact of different combinations and durations of HL and RF on ocular axial elongation, refractive error development, choroidal thickness and other ocular parameters in a chicken model of LIM.

## Methods

### Animals and enclosure

All animals used in this study were treated in accordance with the ARVO Statement for the Use of Animals in Ophthalmic and Vision Research. The study protocol was approved by the Institutional Animal Care and Use Committee (IACUC 2019/SHS/1479) of the Singapore Experimental Medicine Centre (SEMC). The SEMC is accredited by the Association for Assessment and Accreditation of Laboratory Animal Care (AAALAC) International.

One hundred twenty-six one-day-old chicks (mixed Golden Comet/White Leghorn strain) were provided by the National Large Animal Research facility and randomly assigned into 10 groups of 11 to 13 animals each. Animals were raised for 9 days in a 75 cm (length) x 55 cm (width) x 43 cm (height) custom-built enclosure designed to hold two high intensity light-emitting diode (LED) light fixtures. The enclosure walls were fitted with a square wave grating of a repeated sequence of light and dark bars as accommodative cues. The spatial frequency of the stripes ranged between 0.01 to 0.42 cycles/degree depending on the location of the animal within the enclosure. The chicks were fed ad libitum and were raised under a 12/12h light-dark cycle from 7 am to 7 pm. The temperature within the enclosure was maintained at 28 – 32°C. Light and temperature patterns were monitored using a HOBO Pendant data logger (UA-022-64, ONSET, Bourne, MA, USA). At the end of experiment on day 9, chicks were sedated with a mixture of 0.2 ml/kg ketamine and 0.1 ml/kg xylazine and euthanized with an overdose of sodium pentobarbitone to the heart.

### Background and experimental light setup

All chicks were reared under background lighting (150 lux) during the 12/12h light-dark cycle. This was achieved using 6 strips of ultra-bright LEDs (4000K, 2NFLS-NW LED, Super Bright LED, Inc, MO, USA) fixed over the enclosure. For HL, four LED panels of 64 LEDs each (4000K, USHIO Lighting, Singapore) were used to deliver an average of 15,000 lux when measured in different angles of gazes in the enclosure (up, down, left, right, front, back) (Supplementary Figure 1). The light fixtures were controlled using a programmable Helvar DIGIDIM 910 router (Helvar, Dartford Kent, United Kingdom). Light levels and spectra were measured using a calibrated radiometer and spectroradiometer (ILT5000 and ILT950, International Light Technologies, Peabody, MA, USA).

### Myopia induction

Monocular myopia was induced in all chicks from day 1 post-hatching (D1) until D8 using custom-built concave defocusing lenses (Power: -10 ± 0.5 D, total diameter: 12.5 mm, optic zone diameter: 10 mm, base curve: 6.68 mm, La SER Eye Jewelry, FL, USA). The lens was fitted randomly to one eye of the chick using a custom-built 3-D printed lens holder. The lens holder can be clipped on or off a separate base piece that was glued to the down surrounding the chick’s eye. These lens holders ensured the secure positioning of the lenses on the eyes of the animals and allowed for easy and fast removal during cleaning and light exposure. Lenses were worn for 8 days and were cleaned 3 times/day to ensure their optical clarity. The fellow eye was kept uncovered and used as within-animal control.

### Experimental groups

All 10 groups were subjected to monocular LIM. Nine groups were subjected to different interventions including either transient HL (15,000 Lux), RF, or a combination of HL and RF for one of four durations/day (0, 2, 4, or 6h) centered at 12:00 pm. Details on experimental interventions are provided below and in Table 1.

**Table 1:**
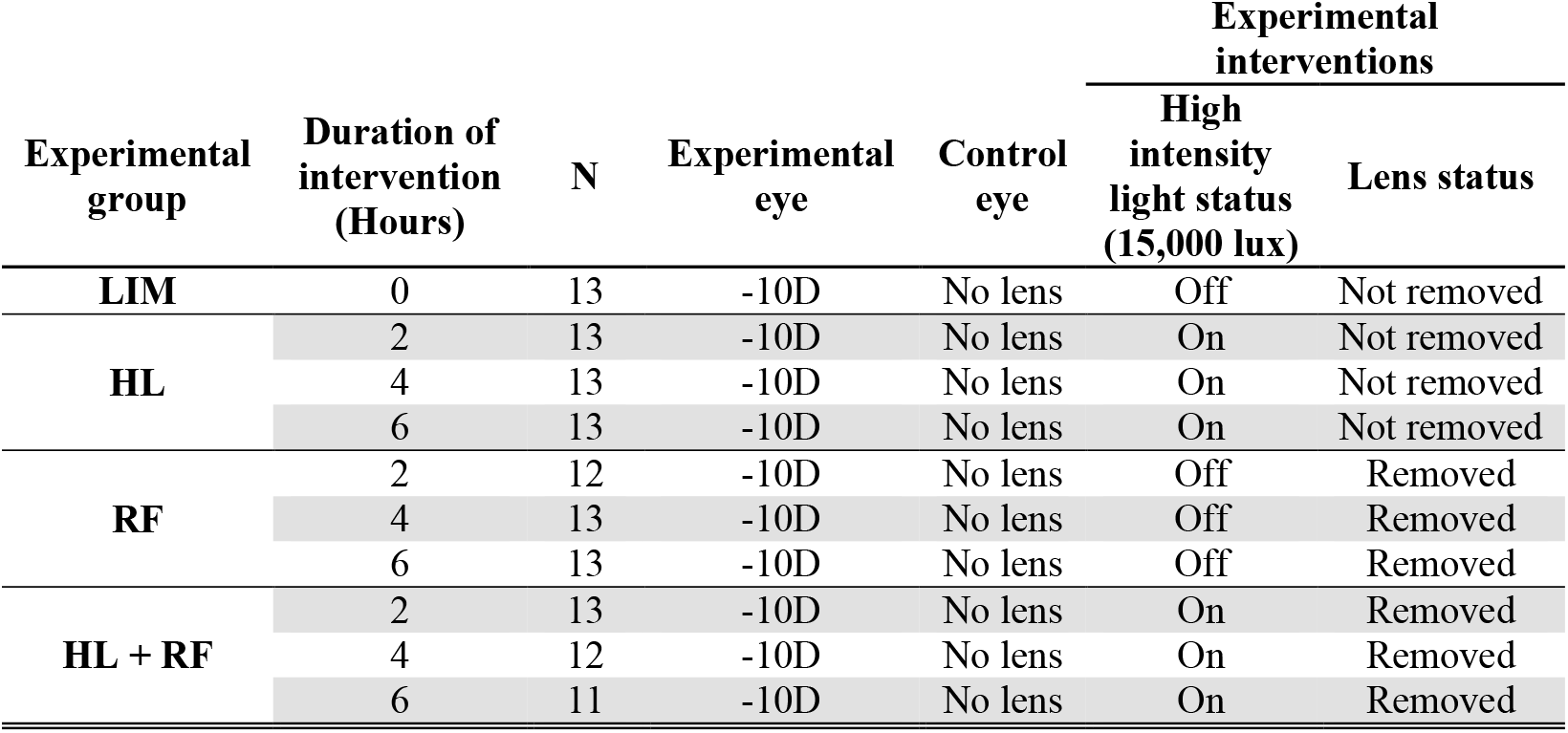
Details on experimental groups and interventions.

#### LIM group

Chicks in this group (*n = 13*) were raised under background laboratory light conditions and were not subjected to HL or RF.

#### High intensity light groups (LIM + HL)

Three groups (*n = 13 each*) were exposed to either 2, 4, or 6h of HL (15,000 lux)/day and background light for the remainder of the light cycle. Defocusing lenses were not removed during HL.

#### Optical re-focus groups (LIM + RF)

Three groups (*n = 13, 12, and 11*) were raised under background light during the light cycle and throughout the experiment. Defocusing lenses were removed for 2, 4 or 6h/day.

#### High intensity light and optical refocus groups (LIM + HL + RF)

Three groups (*n = 12, 13, and 13*) were exposed to 2, 4, or 6h of HL (15,000 lux)/day and background light for the remainder of the light cycle. Defocusing lenses were removed during HL.

### Ocular measurements *in vivo*

On days one (D1), four (D4), and eight (D8), body weight, ocular AL, refractive error, choroidal thickness (CT), central corneal thickness (CCT), and anterior chamber depth (ACD) were measured in the experimental and control eyes of alert, gently hand-held chicks.

Measurements were taken in a dimly lit room (< 5 lux) on all chicks, in a random order, between 12 pm to 5 pm to reduce any impact of circadian rhythm on outcome measures. A lid retractor was only used in chicks who would not keep their eyelids open.

### Axial Length

AL was measured using VuMAX HD (Sonomed Escalon, NY, USA) A-scan ultrasonography with a probe frequency of 10 MHz as described elsewhere.^32^ AL was measured as the distance between the posterior cornea to the anterior retina. Each recorded measurement was the median of 7 to 10 scans.

### Refraction

Ocular refraction was measured using a calibrated automated infrared photo-retinoscope as previously described.^33^ Alert chicks were hand-held on an adjustable platform ∼1 meter away from the infrared photo-retinoscope. The chick’s head was carefully positioned to ensure optimal focus on the chick’s eye and first Purkinje image. Pupil size was adjusted for each eye. The median of the most hyperopic refraction readings (i.e., resting refraction) with no accommodative changes was calculated from the continuous refraction trace comprising of at least 300 readings over time in each eye.^21,33^

### Choroidal thickness and anterior segment characteristics

CT at the posterior pole was measured using posterior segment optical coherence tomography (OCT) (Spectralis, Heidelberg Engineering, Inc., Heidelberg, Germany) and anterior segment (CCT and ACD) was imaged using anterior segment OCT (RTVue, Optovue, Inc., Fremont, CA, USA) as per the protocols described in Najjar et al.^32^ During both procedures, the alert chick’s head was gently held and aligned with the OCT camera lens such that the infra-red laser beam enters the eye through the center of the pupil. The OCT operator further refined the centration of the pupil and multiple OCT scans were captured. For posterior segment OCT measurement, the centration was within ±100 µm from the horizontal line. Distance between the inner border of the sclera and the outer border of the RPE was defined as the CT. The average of three thickness measurements of the central cornea was defined as the CCT whereas, ACD was defined as the distance between the central most posterior layer of the cornea and the central most anterior layer of the lens. Measurements were done manually by the first author SB who, during measurement sessions, was kept blinded to the eye (LIM or control) and study group (HL, RF, HL+RF) conditions.

### Analyses and statistics

All data are represented as mean ± standard error of the mean (SEM) of the inter-ocular difference (IOD) between the experimental and uncovered control eyes (*i*.*e*., Experimental (LIM) Eye - Control Eye) to account for inter-animal differences in outcome measures, given the mixed breed and large number of animals (n = 126 chicks) used in this study. Two-way repeated measures analysis of variance (2wRM ANOVA) with the factors day, group and the group × day interaction were used to compare IODs in refraction, AL, CT, ACD and CCT. A Holm-Sidak method for pairwise multiple comparisons was performed when the omnibus test for interaction effects between group × day was significant. A two-way ANOVA was used to evaluate the interaction between the type of intervention (HL, RF, HL+RF) and its duration (0, 2, 4 and 6h) on refraction, AL and CT. A Holm-Sidak method for pairwise multiple comparisons was used when the omnibus test reached statistical significance. The significance level for all statistical tests were set at α = 0.05 with Sidak correction for post-hoc pairwise comparisons.

## Results

### Ocular changes associated with LIM

Compared to the uncovered contralateral control eyes, the LIM eyes displayed a myopic shift in refractive error that predominantly occurred within the first 4 days after -10D lens wear (IOD: -7.83 ± 0.48 D and -9.02 ± 0.37 D by D4 and D8, respectively). Concurrently, LIM eyes showed an increase in AL (IOD: +0.18 ± 0.02 mm and +0.36 ± 0.04 mm by D4 and D8, respectively) and a decrease in CT (IOD: -26.35 ± 14.55 µm and -90.27 ± 16.44 µm by D4 and D8, respectively), compared to control eyes (all, P < 0.001) (Figures 1-3, Supplementary Table 1). The CCT and ACD of LIM eyes were not different from control eyes.

**Figure 1:**
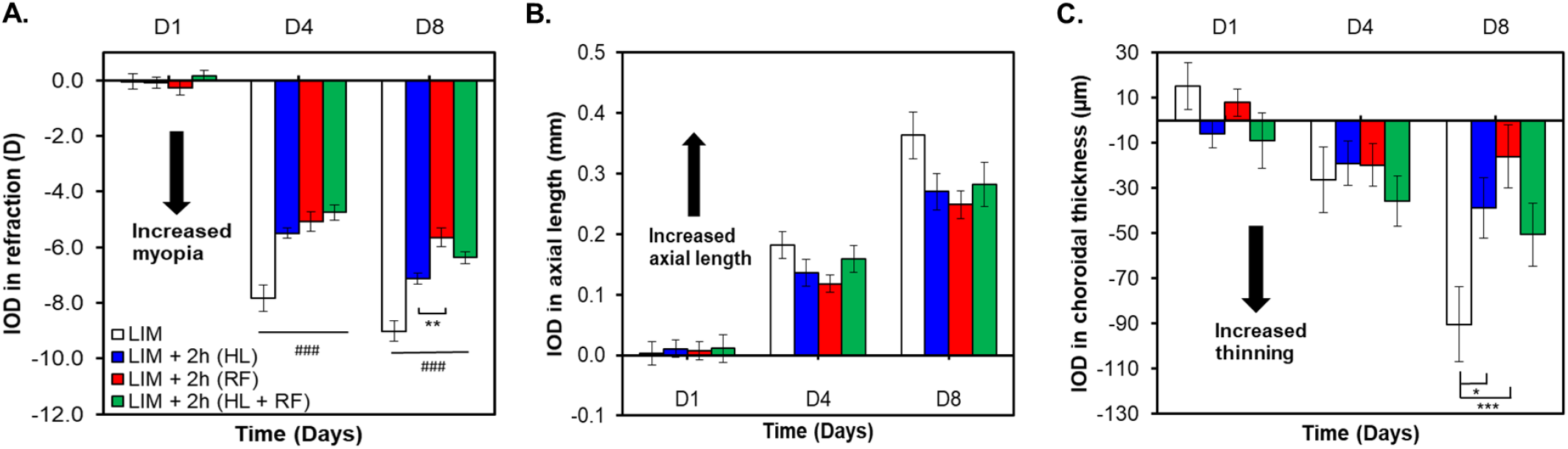
Changes in refraction, axial length and choroidal thickness (expressed as interocular difference (IOD) between LIM eye – control eye) in groups exposed to 0h (LIM) or 2h of HL, RF or both (HL+RF). All groups are significantly different from the LIM group (P<0.001)^###^, (P<0.01)^##^, (P<0.05)^#^ For significant group effect: *: P<0.05, **: P<0.01, ***: P<0.001 (Two-Way RM ANOVA).

### Impact of 2h of HL, RF and HL+RF

For 2h interventions, there was a significant interaction between group and day of intervention for IOD in refraction (F(6, 94) = 10.29, P < 0.001). By D4 and D8, 2h of HL, RF and HL+RF, significantly reduced myopic refraction compared to the LIM group (both for D4 and D8, P < 0.001). Two hours of RF was however more efficient than 2h of HL in reducing myopic refraction on D8 (P = 0.001). HL+RF was as effective as HL and RF in reducing myopic refraction on D4 and D8 (P > 0.05) (Figure 1A). IOD in AL was only dependent on the day of the intervention (F(2, 47) = 161.3, P < 0.001). Differences between groups did not reach statistical significance (Figure 1B). The group × day interaction for IOD in CT was significant (F(6,94) = 3.61, P = 0.003) with RF (P < 0.001) and HL (P = 0.01) significantly reducing choroidal thinning on D8 but not on D4 (Figure 1C). The detailed results are provided in Supplementary Table 1.

### Impact of 4h of HL, RF and HL+RF

For 4h interventions, we found a significant group × day interaction for IOD in refraction (F(6, 94) = 12.80, P < 0.001). By D4 and D8, 4h of HL, RF, and HL+RF, significantly reduced myopic refraction compared to the LIM group (both for D4 and D8, P < 0.001). On D4 and D8, 4h of RF was more effective than HL (P < 0.001) in reducing myopic refraction, while HL+RF was more effective than HL in reducing myopic refraction (D4: P = 0.009, D8: P = 0.04). On D4 but not on D8, RF was more effective than HL+RF (P = 0.01) in reducing myopic refraction (Figure 2A). A significant group × day interaction was also found for IOD in AL (F(6, 94) = 2.23, P = 0.047) with all groups showing significantly reduced axial elongation compared to that observed in the LIM group (D4: no difference between groups; D8: LIM vs RF: P < 0.001; LIM vs HL: P = 0.01; LIM vs HL+RF: P < 0.001) (Figure 2B).

**Figure 2:**
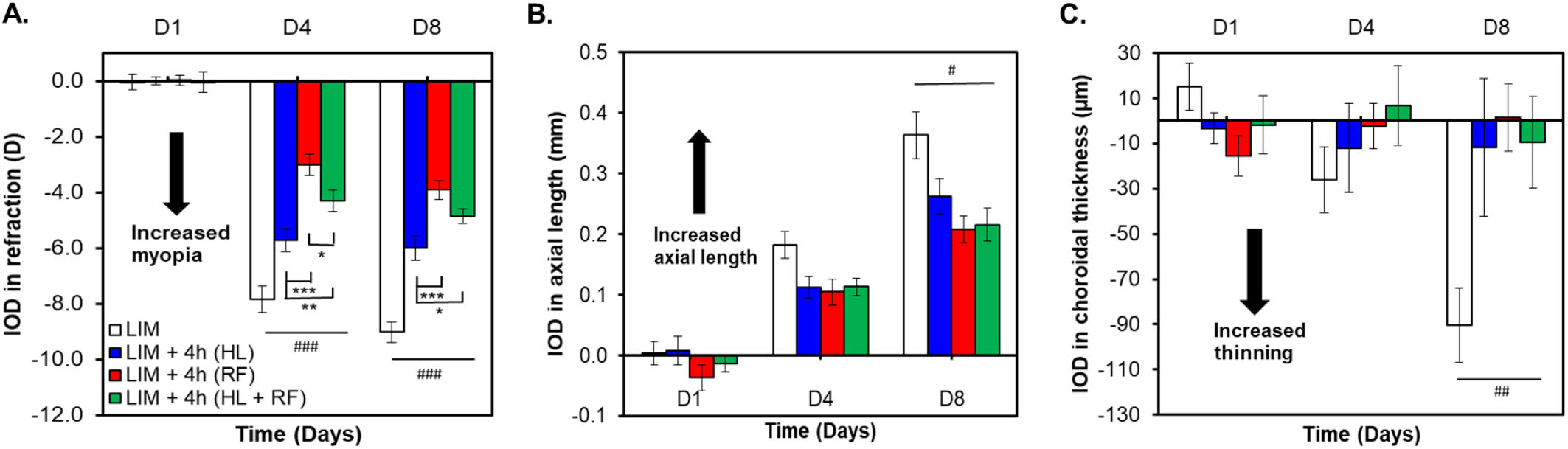
Changes in refraction, axial length and choroidal thickness (expressed as interocular difference (IOD) between LIM eye – control eye) in groups exposed to 0h (LIM) or 4h of HL, RF or both (HL+RF). All groups are significantly different from the LIM group (P<0.001)^###^, (P<0.01)^##^, (P<0.05)^#^ For significant group effect: *: P<0.05, **: P<0.01, ***: P<0.001 (Two-Way RM ANOVA).

Differences in AL between intervention groups did not reach statistical significance. The group × day interaction for IOD in CT was significant (F(6,94) = 3.98, P = 0.001). On D4, increased choroidal thinning was observed in the HL group compared to the HL+RF group (P = 0.049). On D8, all intervention groups prevented choroidal thinning in the myopic eye compared to the LIM group on D8 (LIM vs RF: P < 0.001; LIM vs HL: P = 0.002; LIM vs HL+RF: P = 1.1) (Figure 2C). The detailed results are provided in Supplementary Table 1.

### Impact of 6h of HL, RF and HL+RF

Six-hour interventions had a significant interaction between the group and time for IOD in refraction (F(6,92) = 24.67, P < 0.001). By D4 and D8, 6h of HL, RF, and HL+RF significantly reduced myopic refraction compared to the LIM group (both D4 and D8, P <0.001). Six hours of RF was more effective than HL in reducing myopia refraction by both D4 and D8 (P<0.001). HL+RF was more effective in reducing myopic refraction than both HL (both D4 and D8: P<0.001) and RF (only D8: P = 0.008) (Figure 3A). IOD in AL had a significant group × time interaction (F(6,92) = 6.4, P < 0.001) with all groups showing reduced axial elongation compared to the LIM eyes (LIM vs RF: P < 0.001; LIM vs HL: P = 0.004; LIM vs HL+RF: P < 0.001) on D8. On D4, only the HL+RF group showed significantly reduced axial elongation compared with LIM group (P = 0.009). HL+RF was more effective than HL in reducing axial elongation (P < 0.001) on D8 but not on D4. All interventions (i.e., HL, RF, HL+RF) had similar efficacy in reducing the myopic refraction (P > 0.05) (Figure 3B). CT also showed a significant interaction between groups and time (F(6,92) = 3.53, P = 0.003), with all three interventions significantly reducing choroidal thinning of the myopic eye compared to the LIM group on D8 (LIM vs all groups: P < 0.001) (Figure 3C). Differences in CT between intervention groups did not reach statistical significance.

**Figure 3:**
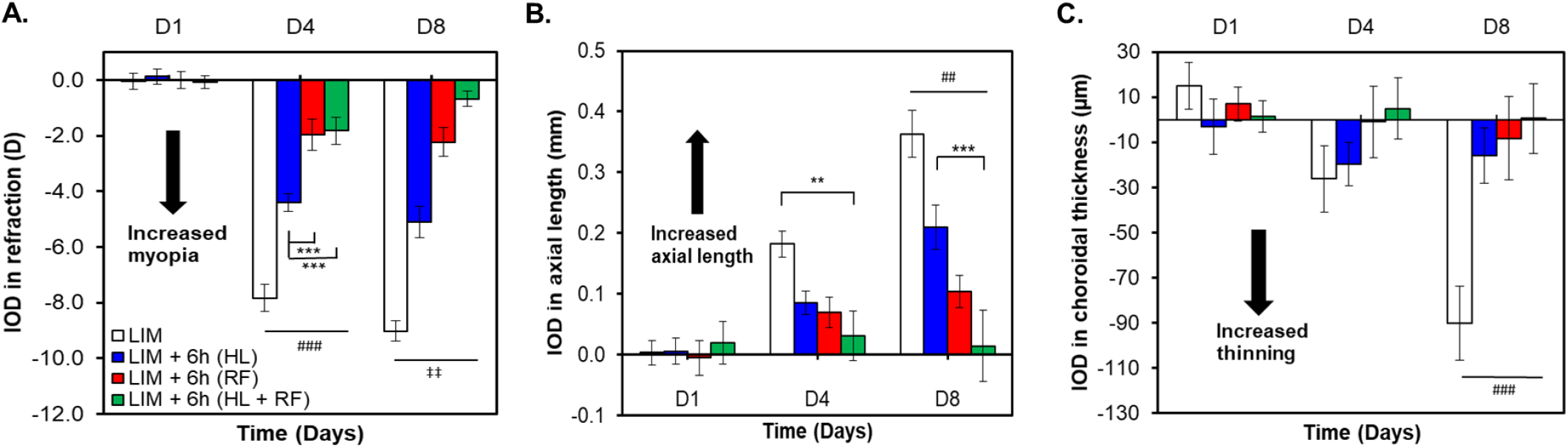
Changes in refraction, axial length and choroidal thickness (expressed as interocular difference (IOD) between LIM eye – control eye) in groups exposed to 0h (LIM) or 6h of HL, RF or both (HL+RF). All groups are significantly different from the LIM group (P<0.001)^###^, (P<0.01)^##^, (P<0.05)^#^ All groups are significantly different from each other (P<0.001)^‡‡‡^, (P<0.01)^‡‡^, (P<0.05)^‡^ For significant group effect: *: P<0.05, **: P<0.01, ***: P<0.001 (Two-Way RM ANOVA).

### Impact of experimental interventions on ACD and CCT

Although ACD showed a significant effect of day for 2h and 6h of any intervention, both ACD and CCT presented with a non-significant change across the intervention groups or their interactions (Supplementary Figures 2-3, Supplementary Table 1).

### Duration response curves on D8 of the interventions

On D8 of the protocol, the impact of interventions (i.e., HL, RF, HL+RF) on IODs of refraction (F(2,104) = 78.33, P < 0.001) and AL (2,104) = 17.1, P < 0.001) was duration-dependent. There was a significant interaction between the duration and type of intervention on IODs of refraction (F(4,104) = 8.38, P < 0.001), where only 6h HL+RF, but not 2h or 4h of HL + RF, was more effective than RF (P = 0.004) and HL (P < 0.001) in reducing myopic refraction. Conversely, regardless of the duration, RF was more effective than HL in reducing myopic refraction (2h: P = 0.01, 4h and 6h: P < 0.001) (Figure 4A). Likewise, the interaction between the duration and type of intervention was significant for AL (F(4,104) = 2.60, P = 0.04), where only 6h of HL+RF was more effective than HL (P < 0.001) (Figure 4B). IOD of CT was not different between intervention groups across the different durations (Figure 4C).

**Figure 4:**
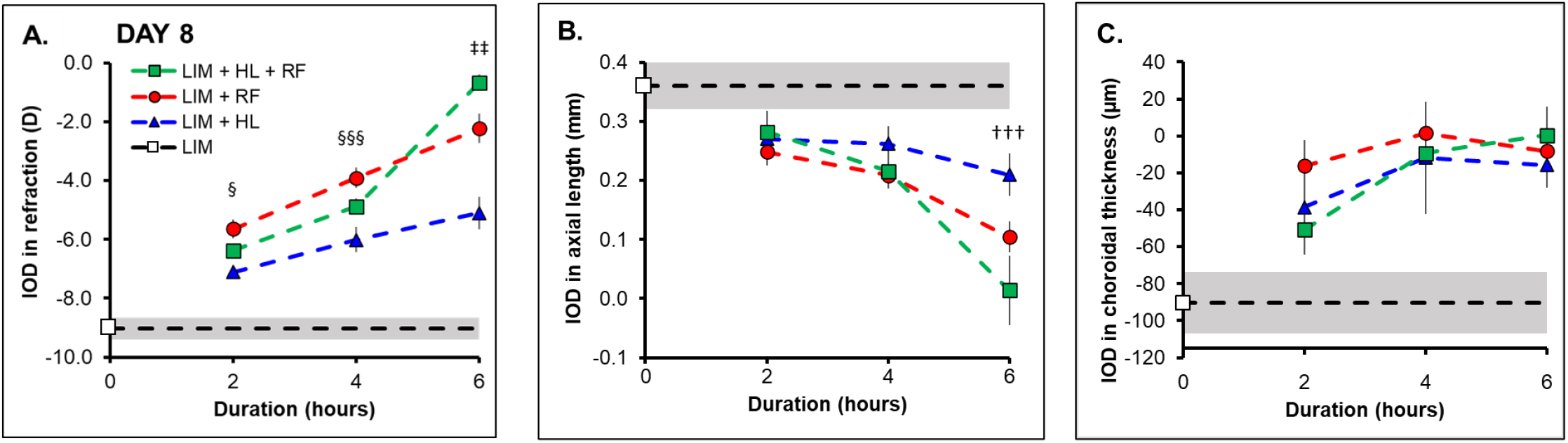
Duration-response curve for refraction (A), axial length (B) and choroidal thickness (C) (expressed as interocular difference (IOD) between LIM eye – control eye) in groups exposed to 0h (LIH group ± 95%CI: white square and shaded area), 2h, 4h and 6h of HL, RF or both (HL+RF) on day 8. HL is significantly different from RF (P<0.05)^§^, (P<0.001)^§§§^ All groups are different from each other (at least, P<0.01)^‡‡^ HL is significantly different from HL+RF (P<0.001)^†††^

## Discussion

In this study, we demonstrated that different combinations and durations of HL and RF have varying effects on the development of LIM in a chicken model. The effect of HL alone, RF alone, and HL+RF in reducing myopia and AL elongation were duration dependent (IODs for refraction and AL were lowest at 6h, followed by 4h, then 2h). We also observed that RF alone was more effective than HL alone in preventing LIM. Interestingly, the effect of HL+RF in reducing LIM development was only found to be additive to a statistically significant extent (lower IODs than both RF alone and HL alone) in the 6h exposure group. On D8, the greatest reduction in LIM (closest to null IODs for myopia, AL, and CT) was found in chicks subjected to 6h daily exposure of HL+RF. We did not find any significant change in ACD or CCT among the groups which was consistent with previous findings.^23^

The duration-dependent effect of RF in reducing LIM development has been previously reported in guinea pigs,^34^ chickens,^35^ tree shrews,^26^ and marmosets.^27^. In chickens, using younger (D1-10) and older (D7-11) animals, Schmid & Wildsoet^35^ reported a ∼90% reduction in LIM (-10 D) by just 3h of normal vision or RF/day. Contrarily, 3h and 6h of RF on older chicks (D7-11) without accommodative aptitudes (ciliary nerve section) reduced their LIM by 60% and 96% respectively, beyond which (i.e., 9h RF) the effect plateaued. In comparison, we observed 37%, 57%, and 75% reduction in LIM by D8 on exposure to 2, 4, and 6h of RF, respectively. The reduced impact of RF that we report in our study may be due to experimental protocol differences, particularly differences in the age (visual maturation) and strain of chickens, the timing of RF (i.e., centered at noon), experimental protocol duration, and the characteristics of the background lighting and visual-spatial environments during normal vision/RF. In addition, Nickla et al showed less ocular growth and LIM during the afternoon than in the morning in response to -10D defocus in chicks.^36^ The authors also found RF to be more effective to reduce LIM in the evening than in the morning.^37^ From our findings, it can be argued that, when centered at noon, longer durations of continuous RF, spilling further into the afternoon (i.e., 2h: 11am – 1pm; 4h: 10am – 2pm; 6h: 9am – 3pm) would potentiate the impact of RF further. Conversely, here we show a newfound duration-dependent impact of HL on reducing myopic shift and axial elongation in LIM. There is currently insufficient evidence linking the total quantum of light exposure (intensity of the exposure × daily duration of the exposure × duration of the study protocol) with LIM reduction, however, a recent study investigating LIM in mice showed that retinal expression of dopamine-related genes and proteins increased with increasing intensities of light,^31^ suggesting a dose-dependent release of ocular growth neuromodulators like dopamine and/or nitric oxide (NO).^38,39^ Whether this release is dependent on the total quantum of light, whereby the retina acts as a photon counter to drive neuromodulatory responses remains unclear yet unlikely, given that short intermittent light pulses have been reported to be more effective than equiluminant continuous light in chickens.^40^ In humans, myopia progression rates in children are slower during a 6-month period that included summer vacation, when days are longer and brighter, compared to winter. These findings partially support an association between the total quantum of light exposure and myopia control.^41^

Next, RF alone was more effective than HL alone in reducing LIM development. During RF, a temporary myopic defocus occurs, while in HL, the hyperopic defocus is constantly present in eyes with LIM. Studies had shown that when the eye receives competing defocus signals, the more myopic defocus dominates ocular growth control;^42–44^ this could explain the stronger emmetropization signal from RF. Another suggestion for the disparity involves RF being a visually/optically guided feedback process,^14,42^ with an endpoint of refractive error elimination. In contrast, HL acting through a potentially distinct pathway involves photoreceptor stimulation, pupillary constriction and retinal neuromodulation^19,22,23^ that can only marginally overcome the existing visual experience (i.e., hyperopic defocus).

Whether RF remains a more potent driver for emmetropization if the myopiagenic stimulus is reduced (e.g., using a -5D defocus instead of -10D), remains to be investigated. Additionally, axial myopic retinas are more prolate (flatter in periphery) and have a relative peripheral hyperopia compared to hyperopic eyes with oblate retinas (steeper in periphery) and relative peripheral myopia.^45,46^ Relative peripheral refraction (hyperopia or myopia) compared to the central refraction may exacerbate or reduce myopia development, respectively,^45,46^ and can partially explain a greater drive of RF alone in reducing LIM compared to HL alone.

Our data did not demonstrate that the combined effect of HL+RF in reducing LIM development was additive, when interventions were 2h or 4h long. These findings suggested that RF and HL may be deploying different pathways of compensatory mechanisms in LIM.^18^ In chicks, Ashby et al.^22^ found that on D5, the protective effects of diffuser removal against FDM was significantly enhanced by 15 minutes of high-intensity indoor lights (15,000 lux) during diffuser removal. This was not in agreement with our findings, where at both D4 and D8, the protective impact of HL+RF on AL increase, myopic refraction, and choroidal thinning, was similar if not worse than RF alone for 2 and 4h-long exposures. The additional alleviation from FDM observed by Ashby and colleagues may result from higher exposure levels to HL and dopamine synthesis due to diffuser removal,^29–31^ while in LIM there was only a minimal increase in HL levels reaching the eye associated with RF. In addition, Ashby et al.^22^ reported that the spectral distribution of their experimental halogen lights (300–1000 nm, peaking at 700 nm) was similar to daylight over the range of visible light for chickens (360–700 nm),^47^ while our experimental light had a typical LED spectrum peaking at 449 nm and 583 nm (Supplementary Figure 1). The fullness of the halogen light spectrum light, may have promoted the impact of short-duration high-intensity light on the recovery from FDM.^21^ Parenthetically, studies investigating the impact of lens or diffuser removal should take into account the different spatial characteristics of the housing environment.^11,48^ Another plausible explanation for our findings was that the significant increase in pupillary constriction induced by HL, during RF, increased the depth of focus and attenuated the accuracy of the RF drive. Pupillary constriction under HL may also explain the reduced impact of defocus on LIM, but failed to explain the potentiating effect of HL on lens induced hyperopia.^23^

Interestingly, HL and RF were additive to a statistically significant extent when interventions were 6h long and almost completely compensated for LIM (IOD: -0.68 D), AL (IOD: 0.01 mm) and CT (IOD: 0.46 µm) by D8 (Supplementary Table 1). These findings suggested a duration-dependent synergy/inter-modulation between HL and RF. While we do not have an understanding of the mechanisms to explain these data yet, this duration-dependent synergy may originate from delayed photobiomodulatory processes originating at the retinal level that enhance the drive for emmetropization conferred by RF.^49^ Vitreal DOPAC (a metabolite of DA) levels are known to increase with both the duration^50^ and intensity^30^ of light exposure in chickens, with the circadian rhythm of DA release observed to peak at 12h into the light phase.^51^ DA can trigger the release of other neurotransmitters like NO^38^, reducing LIM either alone or in conjunction. This DA release is shown to depend not only on the retinal luminance, but also the image contrast / visual spatial information,^52^ which gets altered during normal vision or RF. It is possible that the combination of RF and HL for 6h surpassed a retinal DA threshold that enhanced recovery from LIM further, through an overlapping pathway between RF and HL. It is worth highlighting that the trend of HL and RF alone was unaltered throughout the different durations of exposure. Another potential explanation to our findings is a pupillary escape occurring between 4h and 6h of HL exposure, leading to a reduction in a depth of focus and an optical potentiation of RF. This hypothesis remains to be proven.

In existing literature, chicks had shown changes in CT in response to altered retinal image quality, and it had been suggested that this CT is a part of mechanisms regulating ocular growth and emmetropization.^2,53,54^ As expected, in our study, LIM lead to choroidal thinning.^36,37^ While eyes exposed to 4h and 6h of HL, RF or HL+RF, all had significantly thicker choroids compared to LIM eyes, 2h of HL and RF but not HL+RF lead to significantly thicker choroids than LIM eyes corroborating the observed non-additivity of 2h-long HL and RF exposures for AL and refraction. An increase in choroidal thickness was particularly observed when the duration of interventions was increased from 2h to 4h. Under prolonged HL exposure, higher intraocular temperatures could also increase permeability and vasodilation of choroidal vessels; the increased choroidal blood flow and accompanied release of neurotransmitters from the RPE over time would result in choroidal thickening.^53,54^ This observation could have explained the synergy between HL and RF at 6h interventions, however, the additive effect of HL and RF seems to be independent of choroidal thickening induced by HL as 4h of HL yielded similar choroidal thickening as 6h of HL.

Our study has a few limitations. First, we have opted for an environment lacking color, movement and other usual features of a visual environment, for the housing of the chicks. This environment may mimic urban visual environment lacking greenery, fine spatial details and is probably not the most suitable environment for emmetropization or the recovery from LIM.^11^ Second, given the differences in ocular optics, anatomy and physiology between chickens and humans,^55^ it was difficult to directly contextualize and translate our findings to humans. Nevertheless, our work is in agreement with the literature that removing hyperopic defocus reduces myopia development. Concurrently, exposure to HL can slow the development of myopia in a duration-dependent matter regardless of the refractive status of the eye, or concurrent myopiagenic stimuli (e.g., near visual work). A combination of high intensity light exposure and visual breaks (e.g., reducing accommodative lag) may have the highest potential to prevent myopia development, when applied for long durations during daytime (50% of daytime here). Nevertheless, such lengthy interventions may not be implementable in children.

## Conclusion

In this study, we showed that daily exposure to 2, 4 or 6h of HL or RF slows the shift to myopic refraction in chickens in a duration-dependent manner. Combined with RF, only 6h of HL fully halted the development of LIM (i.e., axial elongation, myopic refraction and choroidal thinning). The synergetic effect of HL and RF is dependent on the duration of the intervention. Further molecular work is required to better understand this peculiar synergy between HL and RF, with the potential of translating such findings into pharmacologic interventions or combined light/optics interventions.

## Supplementary Material

**Supplementary table 1:** **Changes in ocular measurements in groups exposed to 0h, 2h, 4h and 6h of high-intensity light, optical refocus or both. Data represented as mean ± SEM of the interocular difference between experimental and control eyes.**

| Ocular parameter | Duration of intervention (hours) | Protocol | Days |  |  | *P-values 2W RM ANOVA |  |  |
| --- | --- | --- | --- | --- | --- | --- | --- | --- |
|  |  |  | D1 | D4 | D8 | Group | Day | Group × Day |
| Refraction (D) | 0 | LIM | -0.05 ± 0.28 | -7.83 ± 0.48 | -9.02 ± 0.37 | - |  |  |
| Axial length (mm) |  |  | 0.00 ± 0.02 | 0.18 ± 0.02 | 0.36 ± 0.04 |  |  |  |
| Choroidal thickness (μm) |  |  | 15.04 ± 10.35 | -26.35 ± 14.55 | -90.27 ± 16.44 |  |  |  |
| ACD (mm) |  |  | 0.01 ± 0.01 | 0.02 ± 0.02 | 0.02 ± 0.02 |  |  |  |
| CCT (μm) |  |  | -1.41 ± 1.48 | 0.08 ± 1.54 | -2.69 ± 1.37 |  |  |  |
| Refraction (D) | 2 | LIM + HL | -0.08 ± 0.20 | -5.49 ± 0.18 | -7.11 ± 0.20 | <0.001 | <0.001 | <0.001 |
|  |  | LIM + RF | -0.27 ± 0.24 | -5.08 ± 0.36 | -5.65 ± 0.33 |  |  |  |
|  |  | LIM + HL + RF | 0.16 ± 0.19 | -4.75 ± 0.28 | -6.37 ± 0.22 |  |  |  |
| Axial length (mm) |  | LIM + HL | 0.01 ± 0.01 | 0.14 ± 0.02 | 0.27 ± 0.03 | 0.092 | <0.001 | 0.196 |
|  |  | LIM + RF | 0.01 ± 0.02 | 0.12 ± 0.01 | 0.25 ± 0.02 |  |  |  |
|  |  | LIM + HL + RF | 0.01 ± 0.02 | 0.16 ± 0.02 | 0.28 ± 0.04 |  |  |  |
| Choroidal thickness (μm) |  | LIM + HL | -6.08 ± 6.14 | -19.04 ± 9.65 | -38.65 ± 13.39 | 0.206 | <0.001 | 0.003 |
|  |  | LIM + RF | 7.73 ± 5.87 | -33.62 ± 8.88 | -16.12 ± 13.94 |  |  |  |
|  |  | LIM + HL + RF | -9.00 ± 12.33 | -35.83 ± 11.04 | -50.67 ± 13.92 |  |  |  |
| ACD (mm) |  | LIM + HL | -0.01 ± 0.01 | 0.01 ± 0.01 | 0.01 ± 0.01 | 0.576 | 0.022 | 0.845 |
|  |  | LIM + RF | 0.00 ± 0.01 | 0.01 ± 0.01 | 0.03 ± 0.01 |  |  |  |
|  |  | LIM + HL + RF | -0.02 ± 0.01 | 0.01 ± 0.01 | 0.01 ± 0.01 |  |  |  |
| CCT (μm) |  | LIM + HL | 2.68 ± 0.86 | 0.17 ± 0.68 | 1.64 ± 0.63 | 0.015 | 0.59 | 0.274 |
|  |  | LIM + RF | -0.53 ± 0.76 | -0.48 ± 0.67 | -0.15 ± 0.40 |  |  |  |
|  |  | LIM + HL + RF | 0.79 ± 1.07 | 0.10 ± 0.74 | 0.04 ± 0.66 |  |  |  |
| Refraction (D) | 4 | LIM + HL | 0.00 ± 0.14 | -5.71 ± 0.41 | -6.01 ± 0.43 | <0.001 | <0.001 | <0.001 |
|  |  | LIM + RF | 0.01 ± 0.18 | -3.01 ± 0.40 | -3.92 ± 0.35 |  |  |  |
|  |  | LIM + HL + RF | -0.05 ± 0.38 | -4.30 ± 0.38 | -4.87 ± 0.26 |  |  |  |
| Axial length (mm) |  | LIM + HL | 0.01 ± 0.02 | 0.11 ± 0.02 | 0.26 ± 0.03 | 0.001 | <0.001 | 0.047 |
|  |  | LIM + RF | -0.04 ± 0.02 | 0.10 ± 0.02 | 0.21 ± 0.02 |  |  |  |
|  |  | LIM + HL + RF | -0.01 ± 0.01 | 0.11 ± 0.01 | 0.22 ± 0.03 |  |  |  |
| Choroidal thickness (μm) |  | LIM + HL | -3.42 ± 6.85 | -12.08 ± 19.77 | -11.73 ± 30.42 | 0.062 | 0.066 | 0.001 |
|  |  | LIM + RF | -15.58 ± 8.95 | -2.33 ± 9.97 | 1.46 ± 14.93 |  |  |  |
|  |  | LIM + HL + RF | -1.92 ± 12.80 | 6.77 ± 17.47 | -9.38 ± 20.20 |  |  |  |
| ACD (mm) |  | LIM + HL | -0.02 ± 0.02 | 0.00 ± 0.01 | 0.00 ± 0.01 | 0.507 | 0.42 | 0.822 |
|  |  | LIM + RF | -0.01 ± 0.01 | 0.02 ± 0.02 | 0.02 ± 0.02 |  |  |  |
|  |  | LIM + HL + RF | 0.02 ± 0.01 | 0.00 ± 0.01 | 0.02 ± 0.02 |  |  |  |

| CCT (μm) |  | LIM + HL | 1.46 ± 1.11 | -0.67 ± 0.68 | 0.77 ± 1.42 | 0.332 | 0.696 | 0.572 |
| --- | --- | --- | --- | --- | --- | --- | --- | --- |
|  |  | LIM + RF | -0.03 ± 1.43 | 1.03 ± 1.92 | -0.67 ± 1.28 |  |  |  |
|  |  | LIM + HL + RF | -0.49 ± 1.14 | -2.06 ± 1.42 | -0.94 ± 1.16 |  |  |  |
| Refraction (D) | 6 | LIM + HL | 0.12 ± 0.28 | -4.41 ± 0.31 | -5.10 ± 0.56 | <0.001 | <0.001 | <0.001 |
|  |  | LIM + RF | 0.00 ± 0.32 | -1.97 ± 0.55 | -2.23 ± 0.51 |  |  |  |
|  |  | LIM + HL + RF | -0.09 ± 0.23 | -1.83 ± 0.49 | -0.68 ± 0.28 |  |  |  |
| Axial length (mm) |  | LIM + HL | 0.01 ± 0.02 | 0.09 ± 0.02 | 0.21 ± 0.04 | <0.001 | <0.001 | <0.001 |
|  |  | LIM + RF | -0.01 ± 0.03 | 0.07 ± 0.03 | 0.10 ± 0.03 |  |  |  |
|  |  | LIM + HL + RF | 0.02 ± 0.03 | 0.03 ± 0.04 | 0.01 ± 0.06 |  |  |  |
| Choroidal thickness (μm) |  | LIM + HL | -3.23 ± 12.24 | -19.85 ± 9.53 | -15.85 ± 12.24 | 0.004 | 0.003 | 0.003 |
|  |  | LIM + RF | 6.91 ± 7.45 | -1.05 ± 15.86 | -8.32 ± 18.42 |  |  |  |
|  |  | LIM + HL + RF | 1.49 ± 6.97 | 4.87 ± 13.47 | 0.46 ± 15.57 |  |  |  |
| ACD (mm) |  | LIM + HL | 0.01 ± 0.01 | 0.03 ± 0.02 | 0.07 ± 0.3 | 0.382 | 0.001 | 0.433 |
|  |  | LIM + RF | -0.02 ± 0.02 | -0.01 ± 0.01 | 0.05 ± 0.02 |  |  |  |
|  |  | LIM + HL + RF | 0.01 ± 0.01 | -0.02 ± 0.01 | 0.04 ± 0.02 |  |  |  |
| CCT (μm) |  | LIM + HL | -1.54 ± 0.97 | -0.38 ± 0.93 | 0.28 ± 1.14 | 0.437 | 0.21 | 0.876 |
|  |  | LIM + RF | -0.21 ± 2.88 | 2.48 ± 2.31 | 0.18 ± 1.86 |  |  |  |
|  |  | LIM + HL + RF | -1.51 ± 1.11 | -0.38 ± 1.41 | -1.49 ± 0.83 |  |  |  |

\*P values were calculated for 2, 4 & 6 hours of intervention with respect to the control (LIM) group without any intervention (0 hour). All values expressed as the mean interocular difference between experimental - control eyes ± SEM, 2W RM ANOVA: Two-way repeated measures analysis of variance, LIM: Lens induced myopia, HL: High intensity light, RF: Optical refocus, ACD: Anterior chamber depth, CCT: Central corneal thickness

## Duration response curves on D4 of the interventions

On D4 of the protocol, the impact of the intervention on IODs of refraction (F(2,104) = 29.33, P < 0.001) and AL (2,104) = 7.95, P < 0.001) were duration-dependent. The interaction between the group and time for IOD in refraction was significant (F(4,104) = 3.69, P = 0.007), where both 4 and 6 hours of RF (both 4h and 6h: P < 0.001) and HL+RF (4h: P = 0.018 and 6h: P < 0.001) was more effective than HL in reducing myopic refraction. RF of 4h but not 6h, reduced myopic refraction more than HL+RF (P = 0.02) (Supplementary Figure 4A). IOD in AL were not different between groups across the different durations of the interventions (Supplementary Figure 4B). The interaction between the group and time for IOD in CT was significant (F(4,104) = 2.58, P = 0.04), where only 4h of RF (P = 0.009) and HL+RF (P = 0.002) are more efficient than HL in reducing the choroidal thinning of the myopic eyes (Supplementary Figure 4C).

**Supplementary Figure 1:**
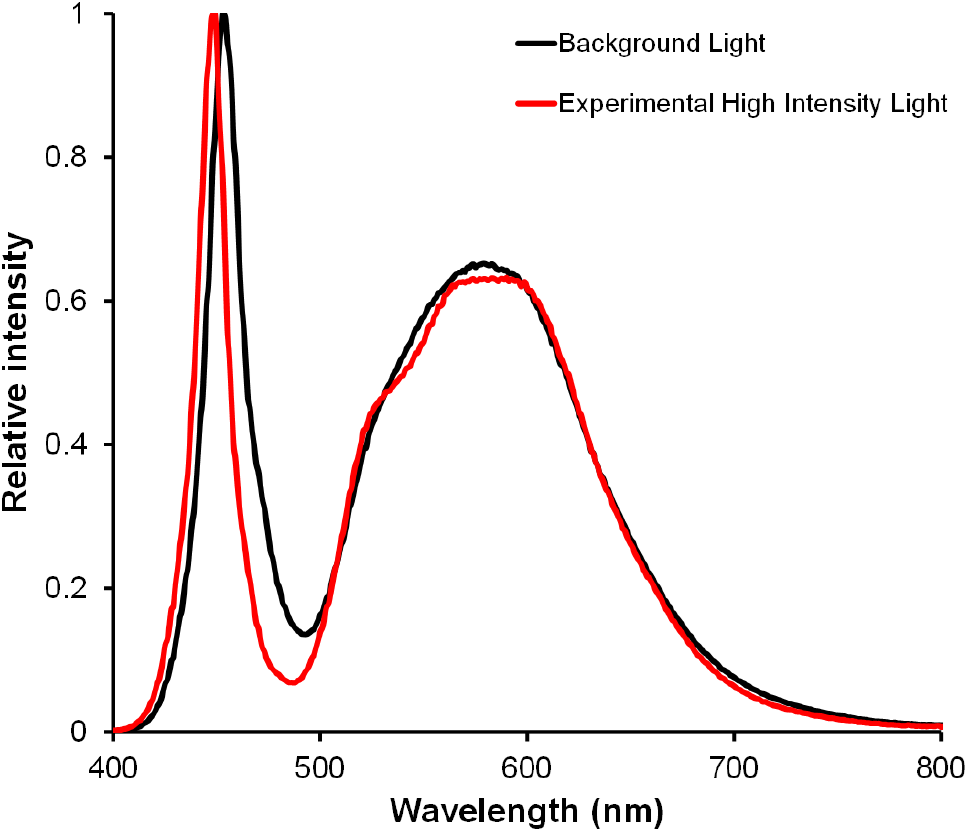
Relative spectral power distribution of the background light. (LED, 4000K, 2NFLS-NW LED, Super Bright LED, Inc, MO, USA), and experimental high intensity light (LED, 4000K, USHIO Lighting, Singapore)

**Supplementary Figure 2:**
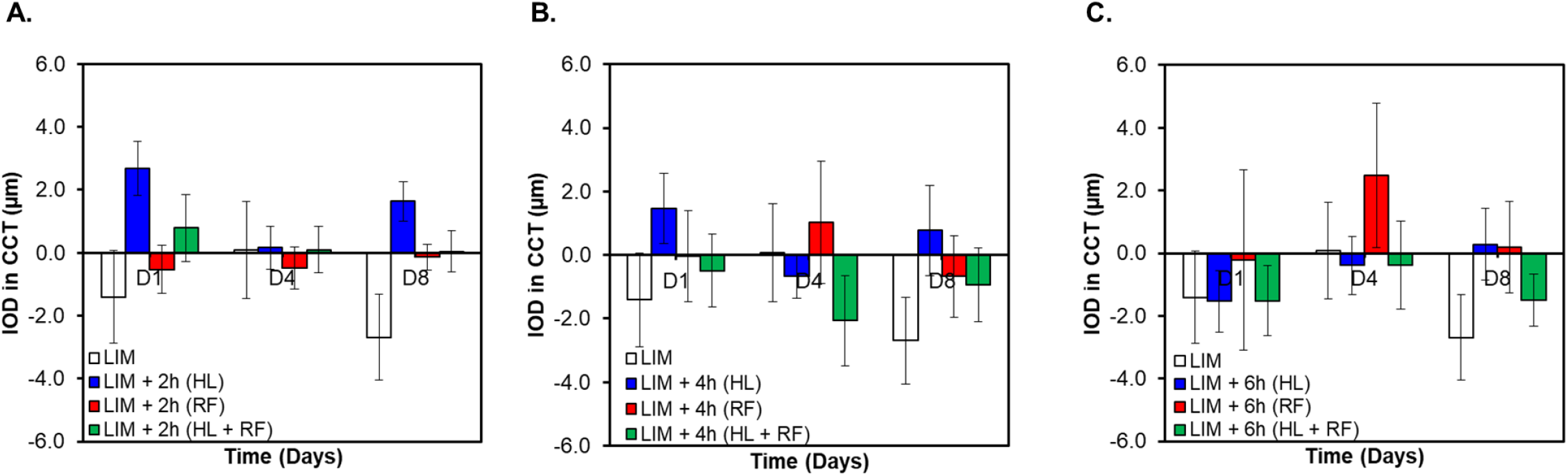
Changes in central corneal thickness or CCT (expressed as interocular difference (IOD) between LIM eye – control eye) in groups exposed to 0h (control), 2h (A), 4h (B) and 6h (C) of HL, RF or both (HL+RF). None of the groups are significantly different from the any other group (Two-Way RM ANOVA).

**Supplementary Figure 3:**
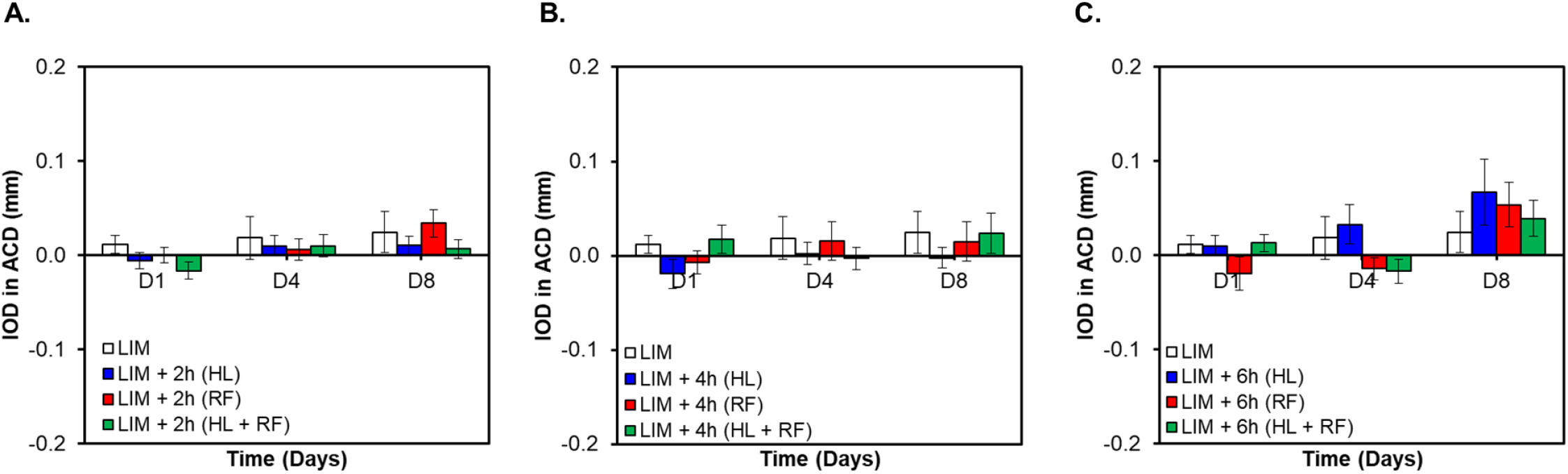
Changes in anterior chamber depth or ACD (expressed as interocular difference (IOD) between LIM eye – control eye) in groups exposed to 0h (control), 2h (A), 4h (B) and 6h (C) of HL, RF or both (HL+RF). None of the groups are significantly different from the any other group (Two-Way RM ANOVA).

**Figure 4:**
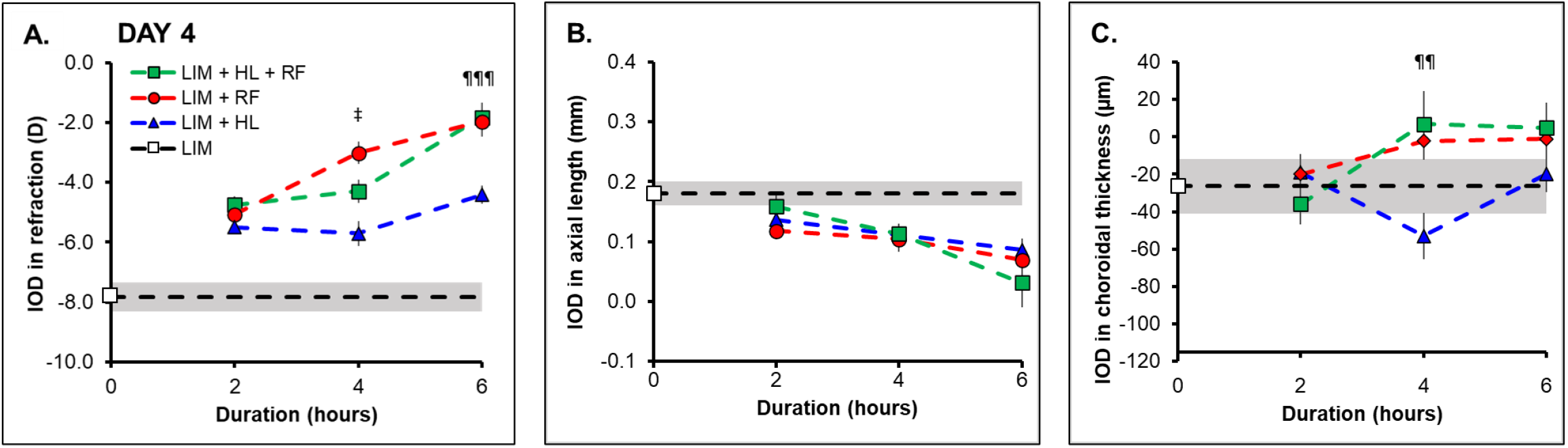
Duration-response curve for refraction (A), axial length (B) and choroidal thickness (C) (expressed as interocular difference (IOD) between LIM eye – control eye) in groups exposed to 0h (LIH group ± 95%CI: white square and shaded area), 2h, 4h and 6h of HL, RF or both (HL+RF) on day 4. All groups are different from each other (P<0.05)^‡^ HL significantly different from both RF and HL+RF (P<0.01)^¶¶^, (P<0.001)^¶¶¶^

